# Chemical composition and larvicidal activity against *Aedes* mosquitoes of flower extracts from *Clitoria Ternatea*

**DOI:** 10.1101/2020.03.31.018465

**Authors:** Darvin R. Ravindran, Madhuri Bharathithasan, Patmani Ramaiah, Mohd Sukhairi Mat Rasat, Dinesh Rajendran, Shakila Srikumar, Intan H. Ishak, Abd Rahman Said, Rajiv Ravi, Mohamad Faiz Mohd Amin

**Affiliations:** Faculty of Medicine, Quest International University, Ipoh, Perak, Malaysia; Royal College of Medicine, Universiti Kuala Lumpur, Ipoh, Perak, Malaysia; Faculty of Earth Science, Universiti Malaysia Kelantan, Jeli Campus, Jeli, Kelantan, Malaysia; School of Biological Sciences, Universiti Sains Malaysia, Minden, Penang, Malaysia; Vector Control Research Unit, School of Biological Sciences, Universiti Sains Malaysia, Minden, Penang, Malaysia; School of Biological Sciences, Faculty of Science and Technology, Quest International University, Ipoh, Perak, Malaysia

## Abstract

**Background:** Mosquito is a human health tread nowadays, the major health problems caused by them are malaria, dengue fever, yellow fever, zika as well as several other outbreaks. The major problem in controlling dengue vectors are the resistant problems. Different classes of insecticides used for public have raised the concern of resistant problem with mosquitoes and environmental pollution. Thus, alternative chemical compounds search is necessary to be searched for overcoming the resistance problem of using pesticides in vectors and the chemical free environment respectively. Thus, to solve these problems, purpose of this study is to identify the larvicidal mechanism, metabolite, antioxidant, chemical compounds and its structure from *C. ternatea* flower and to test its efficacies against early 4^th^ instar larvae of *Ae. aegypti* and *Ae albopictus.*

**Methods:** *Clitoria ternatea* flowers were collected from the Garden of the Faculty of Medicine in International Quest University, Ipoh, Perak and used for crude extraction. Then, the metabolite test, antioxidant test, chromatography techniques were conducted to identify chemical composition of extracts and their chemical structures were identified using GCMS-QP2010 Ultra (Shimadzu). Next, following WHO procedures for larval bioassays, the extracts were used to evaluate against early 4^th^ instar larvae of *Aedes* mosquito vectors.

**Results:** The larvicidal activity of *Clitoria ternatea* flowers extracts evidently affected the early 4^th^ instar larvae of *Aedes* mosquito vectors. The highest larvicidal activity was observed against early 4^th^ instar larvae of *Ae. aegypti* with the LC50 and LC95 values of 1056 and 2491 mg/L, respectively. Meanwhile larvae bioassay test for *Ae.albopictus* recorded with the LC50 and LC95 values of 1425 and 2753 mg/L. Moreover, the results for non target organism test on Guppy fish, *Poecilia reticulate* showed no mortalities with flower extracts at 2500 mg/L and posses no toxic effects on fish. In this study, total of 16 chemical compounds and 6 chemical compounds have been reported to posse’s direct effects on insecticidal, larvicidal and pupicidal effects. Namely 6 chemicals used for insecticidal properties were Glycerin, 2-Hydroxy-gamma-butyrolactone, Neophytadiene, n-Hexadecanoic acid, cis-Vaccenic acid, and Octadecanoic acid with total of 28.7%. *Clitoria ternatea* flower extracts also showed different types of phenols such as anthocyanin, flavonoid and tannin.

**Conclusions:** Our findings showed that crude extract of *Clitoria ternatea* flower bioactive molecules to be effective and may be developed as biolarvicides for *Aedes* mosquito vector control. Furthermore, this study also provided a baseline understanding for future research work on the field applications of *Clitoria ternatea* flower extracts which could be tested for its long term effects on other non target organisms, including human health.

## Background

Mosquito is a human health tread nowadays, the major health problems caused by them are malaria, dengue fever, yellow fever, zika as well as several other outbreaks. Based on World Health Organization [1], western pacific region, northern hemisphere reports on January 2020 dengue updates recorded as follows; Cambodia with higher average 1239 cases weekly compared to period of 2019, Malaysia with lower 4.2% decrease weekly as 2604 cases with 9 death compared to period of 2019, Philippines with 2778 cases with 10 death on 2020, Singapore with 2508 cases, higher compared to period of 2019, Vietnam recorded 1282 weekly cases at 2020. Southern hemisphere, Australia has recorded as 78 dengue cases since beginning of 2020 till 26 February 2020, which is lower compared to period of 2019 [1]. Thus, dengue cases are inevitable in human community and there are many factors contributing towards this global issue.

Currently only physical and chemical methods were commonly used to control mosquito borne diseases. Physical approaches such as mosquito bed nets, mosquito window nets at homes, electrical mosquito rackets are only temperamental solutions [2]. Besides that the use of chemical methods such as temephos and pyrethroids are more prominent with some resistant challenges [2]. Chemical insecticides enhanced biological activity by synthetic chemicals resulting from active or inactive based compounds and explanatory effects of structurally related compounds that counter resistance development which characterizes most single-component bioactive compounds on current mosquitocide classes [2–5]. Additionally, Ravi et al. [4] supported the conceptions of botany control mechanisms for a simple and sustainable method compared to conventional insecticides.

Nothing like current synthetic or chemical insecticides, advantages of plant based insecticides are composed by natural blends of multiple chemical compounds which may act synergistically on both physiological and behavioral processes of mosquitoes [2–6]. Hence, alternative use of bio-insecticides would provide a more accurate solution against Dengue vectors, *Ae. aegypti and Ae.albopictus.*

In line with this safer alternative conception, *Clitoria ternatea* plant has the potential as bio-insecticide to solve the problems of resistance from single based chemical compound. *Clitoria ternatea* is a well-known flower in Asia countries such as Malaysia, Thailand, Philippine, India, China and widely used in Ayurveda and Chinese medicine [7]. Additionally, Malaysians are using the extract of the flower as food colouring and Indians from Kerala region are consuming this plant as vegetable in their daily basis [7]. The potentials of this plant latent in its secondary metabolites such as polyphenol, triterpenoid, flavonoid, glycosides, anthocyanins, tannins and steroid [8]. Till to date, only one study, Mathew et al.[9] has reported the basic use of *C. ternatea* for *Ae. aegypti* and *Ae. stephensi* mosquito larvicidal but none on *Ae. albopictus*. According to Mathew et al. [9], the methanolic extract of *C. ternatea* showed larvicidal effectiveness, however no mechanisms of larvicidal was shown and has suggested future improvements on the investigation of active principles of chemical compounds responsible for larvicidal activities. Thus, within this expansion of framework, this is the first study to elucidate larvicidal mechanism, metabolite, antioxidant and chemical compound identifications from *C. ternatea* flower which may be responsible for *Aedes* larvicidal activities. The objective of this study is to identify the larvicidal mechanism, metabolite, antioxidant, chemical compounds and its structure from *C. ternatea* flower and to test its efficacies against early 4^th^ instar larvae of *Ae. aegypti* and *Ae albopictus.*

## Materials and Methods

### Plant materials

*Clitoria ternatea* was authenticated by hebarium of the University of Malaya and the herbarium voucher number KLU60080. *Clitoria ternatea* flowers (Fig.1) were collected from the Garden of the Faculty of Medicine in International Quest University, Ipoh, Perak. The petals of *C. ternatea* were washed in running tap water to remove adhered debris and soil particle and washed again with dH2O and hot air oven was used to dry the petals at 70⁰C until the petals are dry [10]. The dried petals were grinded into powder by using a laboratory grinder and stored in a vacuum tight container for further usage.

**Fig 1.**
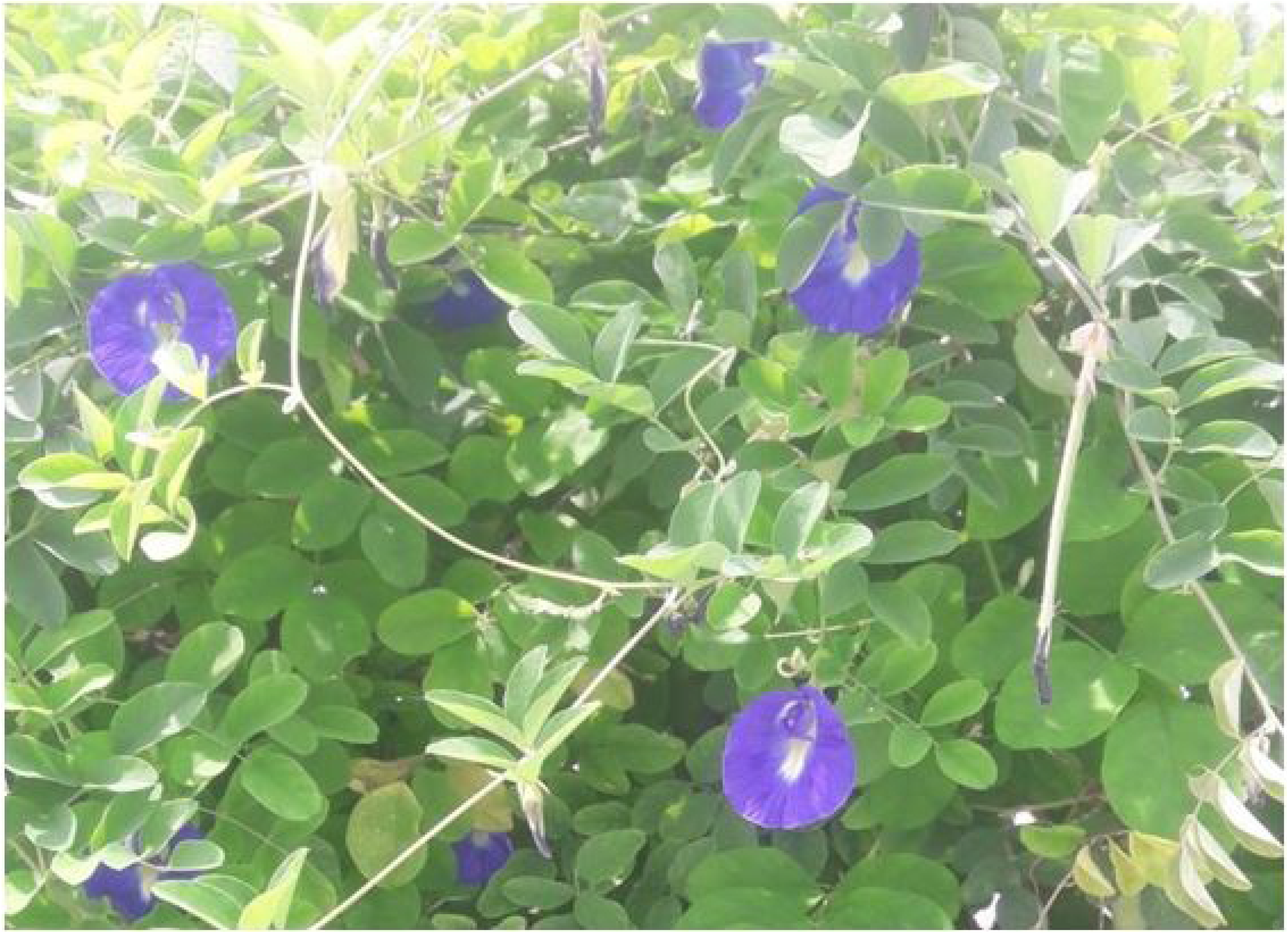
Picture of *Clitoria ternatea* flower from the field

### Aqueous extraction

Aqueous extraction was done by maceration and prepared using 250 ml distilled water and 50 g of powdered *C. ternatea* flowers petal. They were mixed in a 500 ml beaker and stored in a dark area for 1hour. Then, the extraction was filtered using cotton wool and filter paper. At last, the final volumes were stored in a universal bottle and maintain in 4°C to 6°C [10].

### Methanol extraction

Methanol extraction was prepared in three different concentrations; those are 50%, 70%, and 95%. A mixture of 5 g powdered *C. ternatea* flowers petals and 100 ml of respected concentration of methanol was prepared in three separate scott bottle and covered with aluminium foil to avoid light exposure. Later, the bottles were kept overnight in the orbital shaker for maceration process. After 24 hour the extraction was filtered using cotton wool and filter paper by vacuum filtration method. Next, they were concentrated using rotary evaporator with the temperature of 45°C. At last, the remained concentrated amount of extract was stored in a universal bottle and maintain in 4°C to 6°C [4] [11].

### Ethanol extraction

Ethanol extraction was prepared in three different concentrations same as methanol extraction; those are 50%, 70%, and 95%. Maceration process was prepared using 5 g of powdered *C. ternatea* flowers petals and 100 ml of respected concentration of ethanol and mixed in three separate scott bottle, then covered with aluminium foil to avoid light exposure. Later, the bottles were kept in room temperature for 24 hours. The next day all the extracts were filtered using cotton wool and filter paper by vacuum filtration method. Next, they were concentrated using rotary evaporator with the temperature of 60°C. At last, the concentrated extracts were stored in a universal bottle and maintain in 4°C to 6°C [11].

### Petroleum ether extract

Petroleum ether extraction was done with 5 g of fresh flowers; they were cut into small pieces and soaked in a scott bottle with 50 ml of petroleum ether for 48 hours. Later, extraction was filtered out using filter paper and transfer to a universal bottle and stored in 4°C to 6°C [12].

### Ethyl acetate extraction

Ethyl acetate extraction was prepared with 5 g of powdered *C. ternatea* flowers petals and 100 ml of ethyl acetate in a scott bottle, then covered with aluminium foil to avoid light exposure. Then, the bottle was kept in room temperature for 48 hours and after 48 hours the extract was filtered out using cotton wool and filter paper. At last they were stored in a universal bottle and maintain in 4°C to 6°C [13].

### Test for flavonoid

Flavonoid was determined with two different chemical tests for all nine extractions; 1 ml of *C. ternatea* extracts were added into a test tube followed by a few drops of FeCl_3_, the colour changes were observed. The second confirmation test for presence of flavonoid was done in a new test tube by adding 1 ml of crude extract and few drops of 5% of AlCl_3_ solution and the colour changes were observed [14].

### Test for Anthocyanin

Anthocyanin was determined with two different chemical tests for all nine extractions. First, 1 ml of crude extract was added into a test tube and followed by 2 ml of HCl, then heated for 5 min at 100C, and the color changes were observed. Second test, 1 ml of crude extract was added into a test tube and followed by 2 ml of NaOH, and then the color changes were observed [14].

### Test for Tannins

Tannins were determined with a chemical test for all nine extractions; 1 ml of crude extract was added into a test tube and followed by few drops of FeCl_3_. At last the colour changes were observed [14].

### Test for Polyphenol

UV – visible spectrophotometer was used to evaluate the concentration of polyphenols in each extraction, the absorbance was measured with 725 nm. All the measurement was triplicated for each sample [13].

### Paper chromatography

Paper chromatography was used to study the fractions of component. Chromatography paper were used to study the fractions of the extractions by placing 10 μl of crude extract and allow them to dry completely before immersing in the solvent. The solvent used in this process were the similar solvent that used for the extraction process such as aqueous, 50% methanol, 70% methanol 95% methanol, 50% ethanol, 70% ethanol 95% ethanol, petroleum ether and ethyl acetate. At last the Rf value was calculated [15].

### Thin layer chromatography

Thin layer chromatography is also a method to study components based on fractions. The solvent used for thin layer chromatography is a mixture of methanol and chloroform and this process was done in a specific TLC tank. First the TLC gel was spotted by all the crude extract and they were allowed to dry for few minutes and placed the gel into the tank to immerse with the solvent. Lastly 1% FeCl_3_ was sprayed to identify the spot and fix the spot; at last Rf value was calculated based on the measurements [9].

### Antioxidant activity

Based on the Fu et al. [16], antioxidant scavenging activity of *C. ternate* petals extractions were measure using DPPH solvent. To conduct the study a 96 well plate was used to mix the extract with the DPPH solvents and other controls. Later the plate was covered with aluminium foil and left in a dark area for 30 minutes. During this test ascorbic acid was used as the positive control and Tris hcl buffer was used as the negative control; lastly the color changes was observe and OD was collected using the ELISA reader at 517nm. The scavenging activity of DPPH radicals was calculated using the following equation:

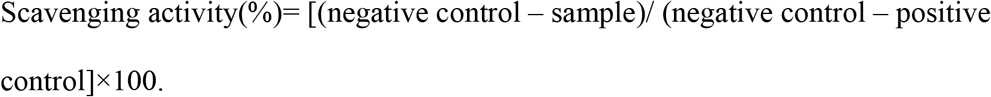

### GC – MS analysis

The GC-MS analysis was only done for methanol extract and was used to determine the compounds, the analysis was performed on a GCMS-QP2010 Ultra Shimadzu. The GCMS-QP2010 Ultra Shimadzu system, fitted with Rtx-5MS capillary column the measurement will be 30 m × 0.25 mm inner diameter, ×0.25 *μ*m film thickness and the maximum temperature can be used is 370°C, coupled with a QP2010 Ultra Shimadzu MS. Ultra-high-purity helium was used as carrier gas at a constant flow rate of 1.0 mL/min. The injection, transfer line, and ion source temperatures were all 280°C. The oven temperature was programmed from 80°C and it was hold for 2 min to 280°C at a rate of 3°C/min. The crude samples were diluted with appropriate solvent and filtered. The particle-free diluted crude extracts about 1 *μ*L were taken in a syringe and injected into an injector with a split ratio of 10:1. All data were obtained by collecting the full-scan mass spectra within the scan range of 40–550 amu. The percentage composition of the crude extract constituents was expressed as the percentage by peak area. The identification and characterization of chemical compounds in various crude extracts were based on the GC retention time [2,4].

### Larvae Rearing

The eggs of *Aedes* were obtained from Vector Control Research Unit (VCRU) at University Sains Malaysia (USM), Penang, Malaysia. We follow the method used by [2–5] in larvae rearing. The eggs were hatched in de-chlorinated water for 24 hours and were maintained at 25 ̊C to 30 ̊C (room temperature), pH of 6.95 to 7.03, and relative humidity of 80 ± 10% and dissolved oxygen from 5.5 to 6.1 mg/L in the laboratory. After five days, the late 3rd instar larvae will be used for bioassay test.

### Larvicidal Bioassay

Larvicidal bioassays were performed in accordance with the standard World Health Organization (2005) [17]. Bioassays were carried out using 25, *Ae. aegypti* and *Ae. albopictus* late 3rd instar larvae (homogeneous population consisting 5 mm to 6 mm in body length). The bioassays were replicated four times using 25 larvae for each concentration with Methanol CH_3_OH as solvent control. During the larvae testing period, fish meal was provided. Initially before selecting the accurate testing dose, all the larvae were subjected to a wide range of test concentrations. This step is necessary to determine the range of extract solution for larvicidal activities [2,4,5]. In this study, seven concentrations ranging from 500mg/L to 2500mg/L, yielding between 0 and 100% mortality in 24 hours of exposure was selected as test concentrations. Control solutions were prepared with 1ml of distilled water and 10% of respective methanol solvent during each experimental replicates [2,4,5]. The reason for using solvent control is to ensure that all test replicates are identical to plant extract solutions and to ensure mortality results were not due to its solvents [2,4,5]. Experiments were conducted at room temperature 28±2°C and larvae mortalities were recorded at intervals of 24 hours and 48 hours [14]. Immobilization and total absence from the larvae, even after touch is the end point of bioassay [2,4,5]. The data were analyzed by using probit analysis in IBM SPSS Statistics 24.

### Morphological view

*Ae. aegypti* early 4^th^ instar larvae was observed under the optical microscope (Leica USA), magnification 40-400x [4].

### Non-targeted organism test

Guppy fish, *Poecilia reticulate*, total of ten fishes in three replicates with 1.20g mean weight and 3.5cm mean length were used in this test (acclimatization period of 12 days in laboratory conditions before the start of experiment). Each treatment group has been tested with larvicidal LC_95_ plant extract concentration dissolved in 2000mL anti-chlorinated water. A control group has also been set with only 2000mL of anti-chlorine treated water. The tests were performed for 24 hours with observations made at first 10 minutes, then 1, 2, 3, 6 and 24 hours. The mortality and observable abnormalities of fish were recorded. Test conditions of water such as pH, water temperature and dissolved oxygen were recorded during the start and end of experiment [4].

## Results

### Metabolite study

Based on the Table 1 and Figure 2 the absorbance showed higher presence of anthocyanin, flavonoid and tannins in the *C. ternatea* flower extracts of 95% methanol compared to other extractions.

**Table 1:**
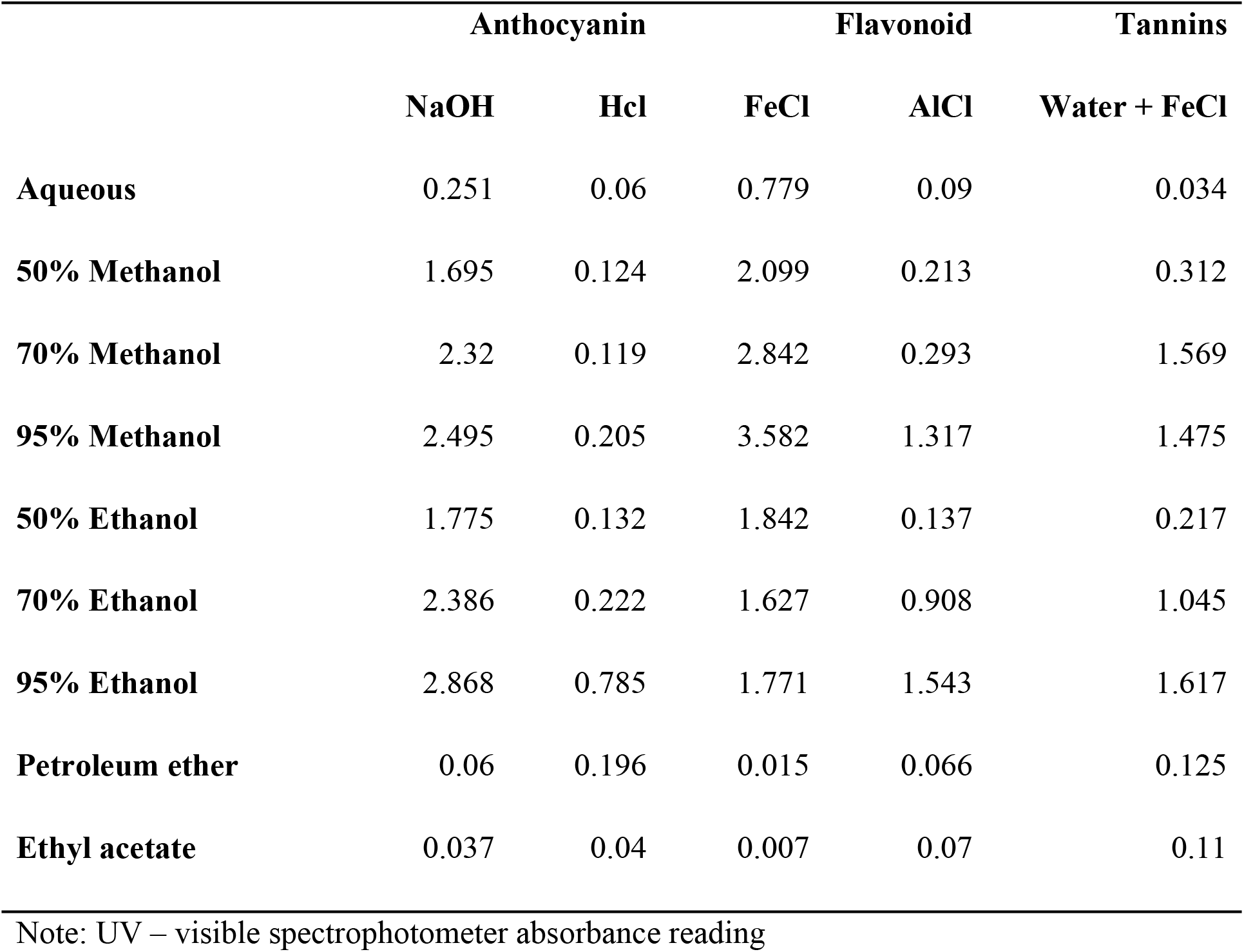
Metabolite study of *Clitoria ternatea* flower extracts

|  | Anthocyanin |  | Flavonoid |  | Tannins |
| --- | --- | --- | --- | --- | --- |
|  | NaOH | Hcl | FeCl | AlCl | Water + FeCl |
| **Aqueous** | 0.251 | 0.06 | 0.779 | 0.09 | 0.034 |
| **50% Methanol** | 1.695 | 0.124 | 2.099 | 0.213 | 0.312 |
| **70% Methanol** | 2.32 | 0.119 | 2.842 | 0.293 | 1.569 |
| **95% Methanol** | 2.495 | 0.205 | 3.582 | 1.317 | 1.475 |
| **50% Ethanol** | 1.775 | 0.132 | 1.842 | 0.137 | 0.217 |
| **70% Ethanol** | 2.386 | 0.222 | 1.627 | 0.908 | 1.045 |
| **95% Ethanol** | 2.868 | 0.785 | 1.771 | 1.543 | 1.617 |
| **Petroleum ether** | 0.06 | 0.196 | 0.015 | 0.066 | 0.125 |
| **Ethyl acetate** | 0.037 | 0.04 | 0.007 | 0.07 | 0.11 |

Note: UV – visible spectrophotometer absorbance reading

**Fig 2:**
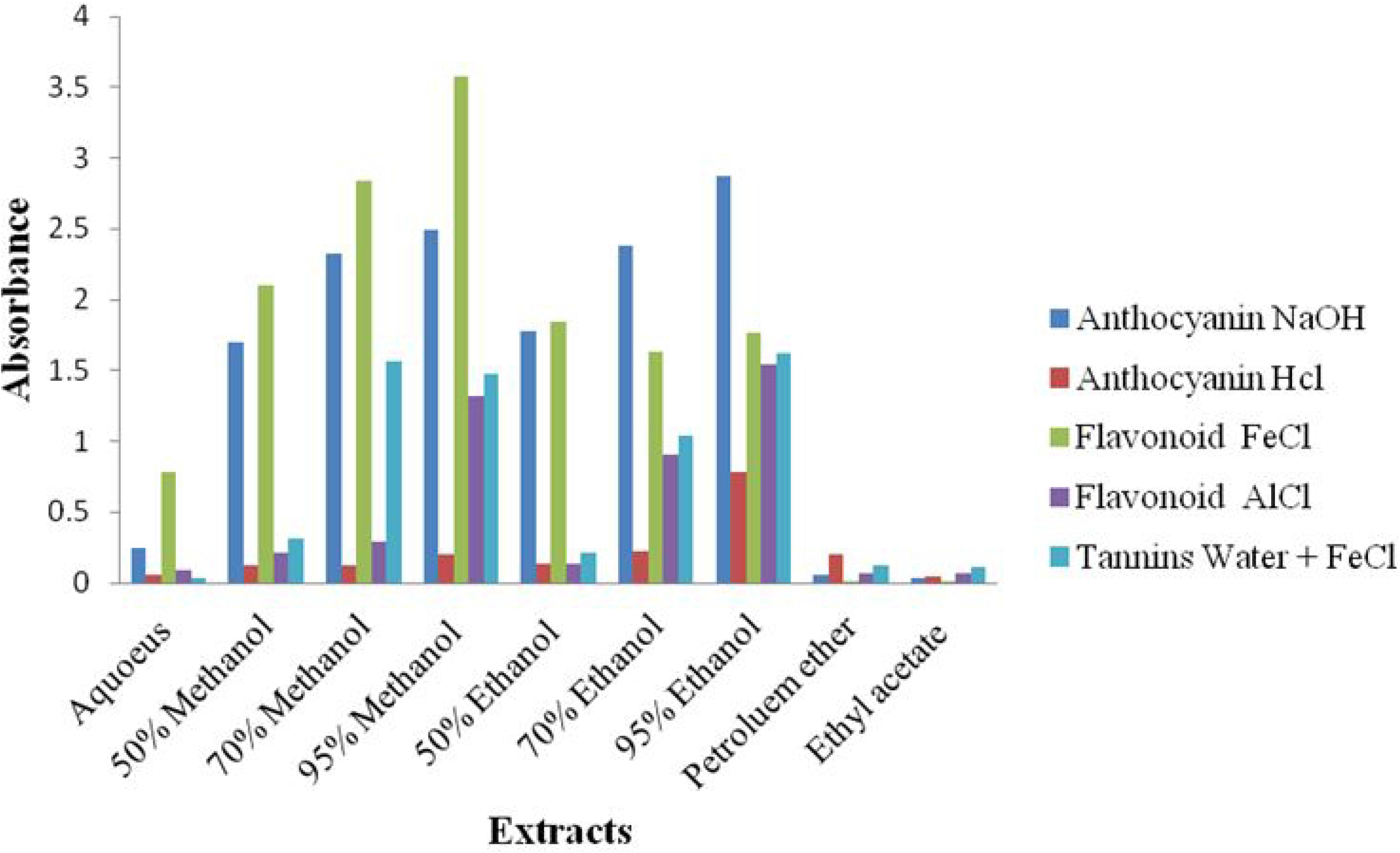
UV – visible spectrophotometer absorbance reading of *Clitoria ternatea* flower extraction for metabolite study.

### Paper chromatography

All the values in Table 2 shows the three Rf values present in *Clitoria ternatea* flowers extracts in all the solvents used for extraction process. The use of methanol solvent showed high Rf values compared to other solvents.

**Table 2:**
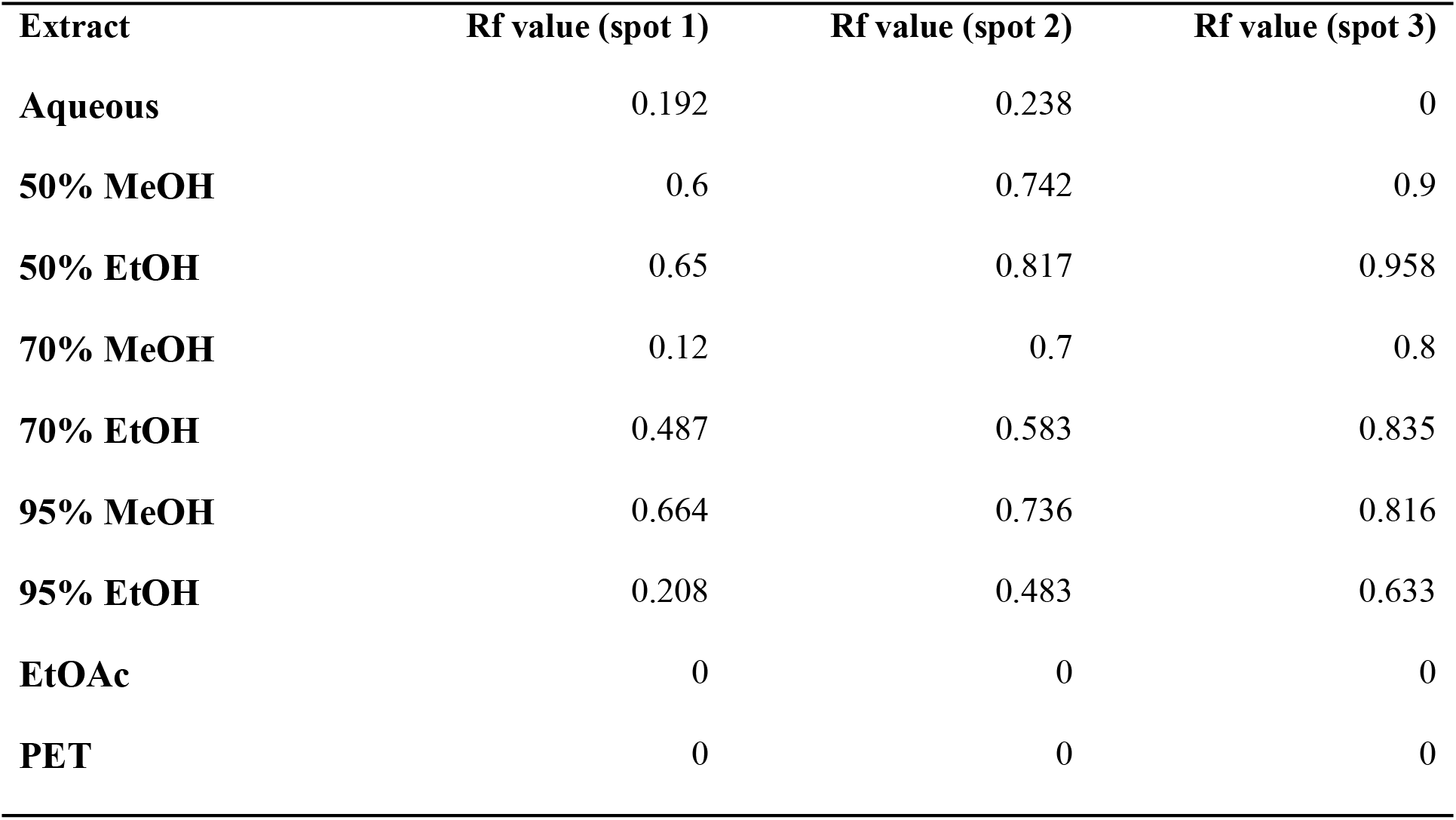
Rf value of paper chromatography for *Clitoria ternatea* flowers extracts.

### Thin layer chromatography

All the values in Table 3 shows the three Rf values present in *C. ternatea* flowers extracts in all the solvents used for extraction process. The use of methanol and ethanol solvent showed approximately same Rf values.

**Table 3:** Rf value of thin layer chromatography *Clitoria ternatea* flowers extracts.

| Extract | Rf value spot 1 | Rf value spot 2 |
| --- | --- | --- |
| Aqueous | 0 | 0 |
| 50% MeOH | 0.55 | 0.639 |
| 50% EtOH | 0.561 | 0.7 |
| 70% MeOH | 0.567 | 0.711 |
| 70% EtOH | 0.544 | 0.644 |
| 95% MeOH | 0.522 | 0.683 |
| 95% EtOH | 0.528 | 0.611 |
| EtOAc | 0 | 0 |
| PET | 0 | 0 |

### Antioxidant activity

2,2-Diphenyl-1-picrylhydrazyl (DPPH) was used to identify the presence of antioxidant property in each extract, based on the Figure 3 the graph shows the extract with high antioxidant property and low antioxidant property. The extract that carries high antioxidant property was aqueous extract.

**Fig 3:**
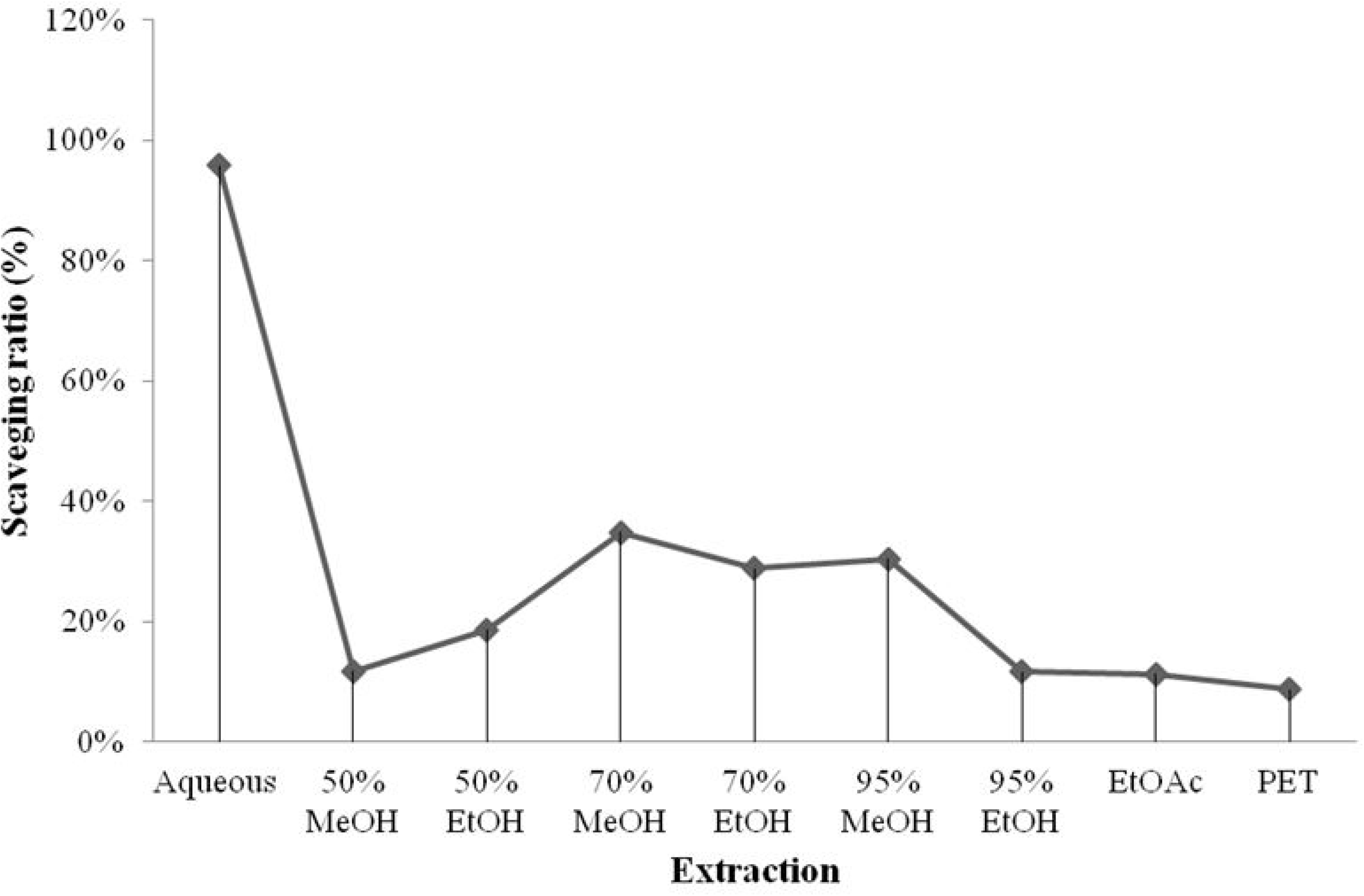
Antioxidant activity of *Clitoria ternatea* flower extracts.

### GC-MS analysis and identification of compounds

GC-MS analysis of methanol solvent extracts of *C. ternatea* flower showed 16 peaks which indicating the presence of 16 phytochemical compounds (Fig 4). On the comparison of the mass spectra of the constituents with the NIST 08 library, 16 compounds were characterized and identified (Tables 4). The major 3 highest peaks chemical compounds in extracts were Acetic acid, 1-(2-methyltetrazol-5-yl)ethenyl ester (16.5%), 4H-Pyran-4-one, 2,3-dihydro-3,5-dihydroxy-6-methy (13.9%) and dl-Glyceraldehyde dimer (12.4%). The attached supplementary pdf file contains the NIST 08 library search for chemical compound structures and details.

**Table 4:**
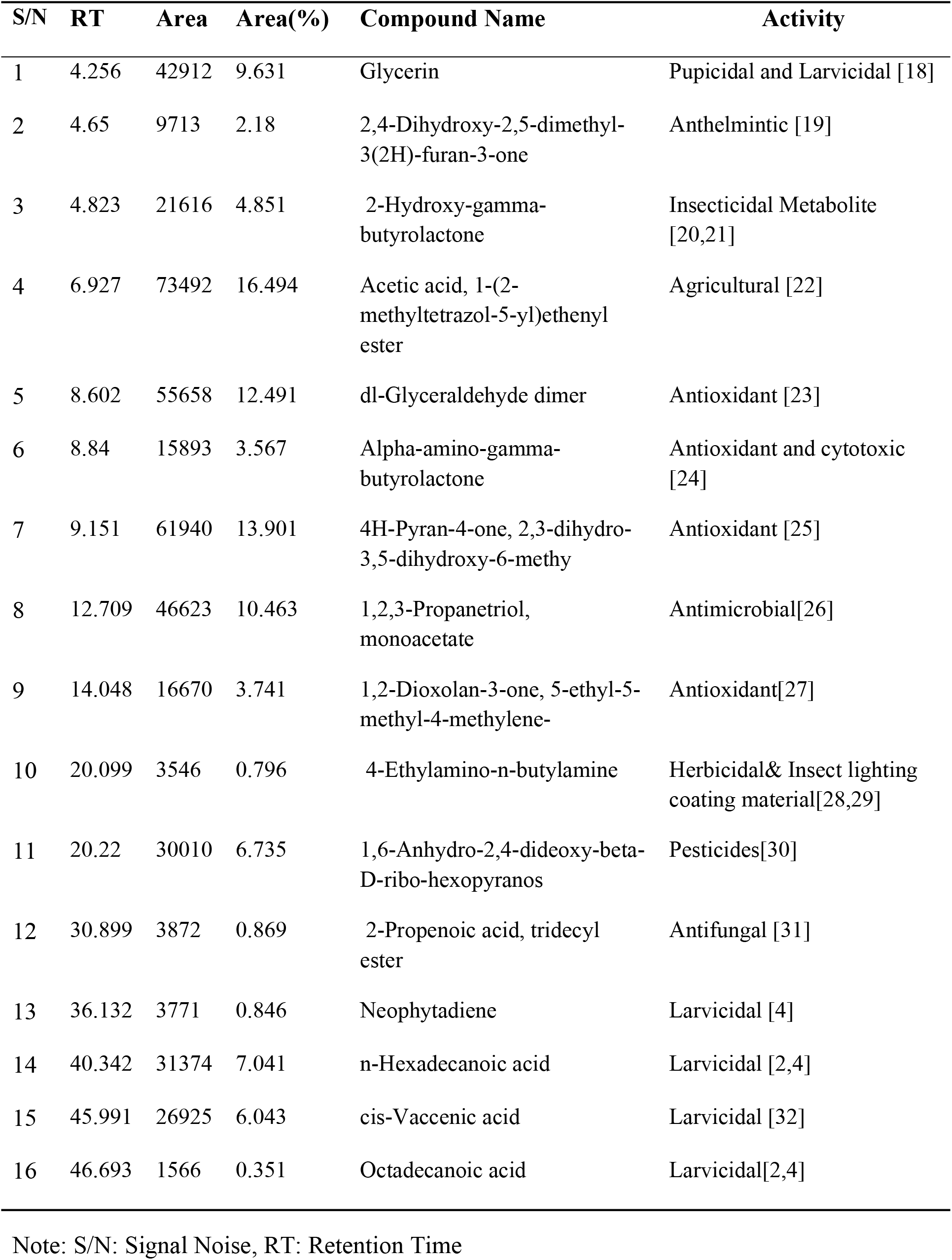
Chemical compounds for *Clitoria ternatea* flower

**Fig 4:**
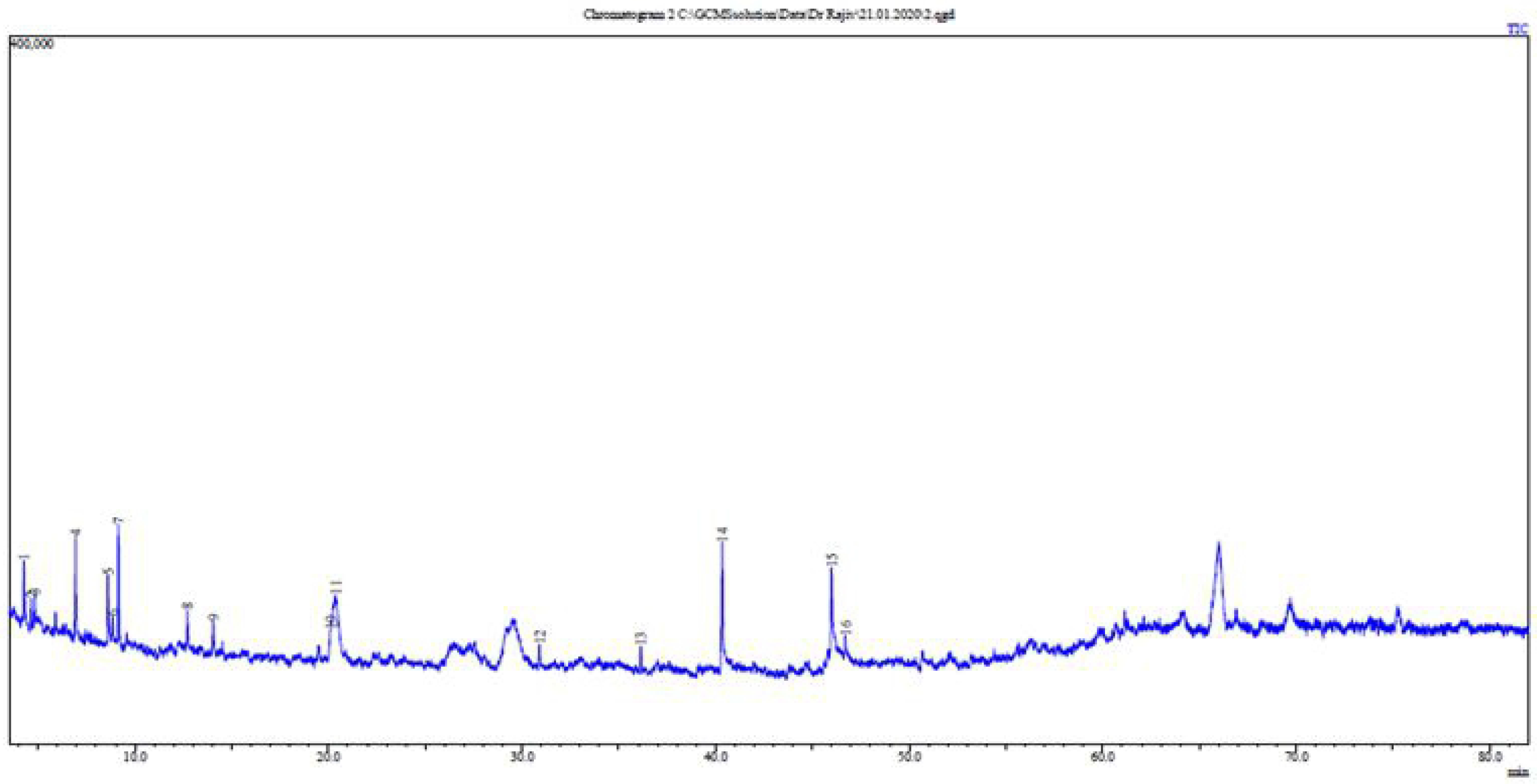
Chromatogram for GC-MS analysis of methanol extract for *Clitoria ternatea* flower extracts (ACQUISITION PARAMETERS; Rtx-5MS capillary column 30 m×0.25 mm inner diameter, ×0.25 μm film thickness, the oven temperature was programmed from 80 °C (hold for 2 min) to 280 °C at a rate of 3 °C/min Carrier Gas = He)

### Larvicidal Bioassay

The bioassay testing from methanol solvent extracts of *C. ternatea* flower were tested at 300 mg/L, 500 mg/L, 1000 mg/L, 1500 mg/L, 1700 mg/L, 2000 mg/L and 2500 mg/L. The entire larvae bioassay test with *C. ternatea* flower extracts showed a significant increase in mortality percentage with the increase of concentration. Among the plant extracts tested, the highest larvicidal activity was observed against early 4^th^ instar larvae of *Ae. aegypti* with the LC50 and LC95 values of 1056 and 2491 mg/L, respectively (Table 5). Meanwhile larvae bioassay test for *Ae.albopictus* recorded with the LC50 and LC95 values of 1425 and 2753 mg/L (Table 5). Figure 6 shows the graphical representation of larvae mortality rate between *Ae. aegypti* and *Ae.albopictus*. Finally, the results for non target organism test on Guppy fish, *Poecilia reticulate* showed no mortalities with plant extracts at 2500 mg/L and posses no toxic effects on fish (Figure 5). The 95% confidence limits LC50 (95%CI) and LC95 (95%CI), chi-square and degree of freedom (df) values were also calculated (Table 5). In control assay there was no any significant mortality.

**Table 5.** Larvicidal activity of *Clitoria ternatea* flower extracts against early 4^th^ instar larvae of *Ae. aegypti* and *Ae. albopictus*

| Larvae | N<sub>a</sub> | LC<sub>50</sub>(mg/L)<br>(95% LCL-UCL) | LC<sub>95</sub>(mg/L)<br>(95% LCL-UCL) | χ<sup>2</sup> | df | R |
| --- | --- | --- | --- | --- | --- | --- |
| *Ae.aegypti* | 100 | 1056<br>(981-1121)<br>Y=-14.142+4.722X | 2491<br>(2094-2641)<br>Y=-14.142+4.722X | 30* | 22 | 0.667 |
| *Ae.albopictus* | 100 | 1425<br>(1352-1491)<br>Y=-19.682+6.282X | 2753<br>(2374-2881)<br>Y=-19.682+6.282X | 30* | 22 | 0.882 |

N<sub>a</sub>; total number of mosquitoes larvae used; n = 25 with 4 replicates, LC50; Lethal concentration 50% mortality, LC95; Lethal concentration 95% mortality, LCL; lower confidence limits, UCL; upper confidence limits, (χ<sup>2</sup>); Pearson chi square, df; degrees of freedom, R; Pearson's R (Note: Chi-square values with asteric are significant P<0.05).

**Fig 5.**
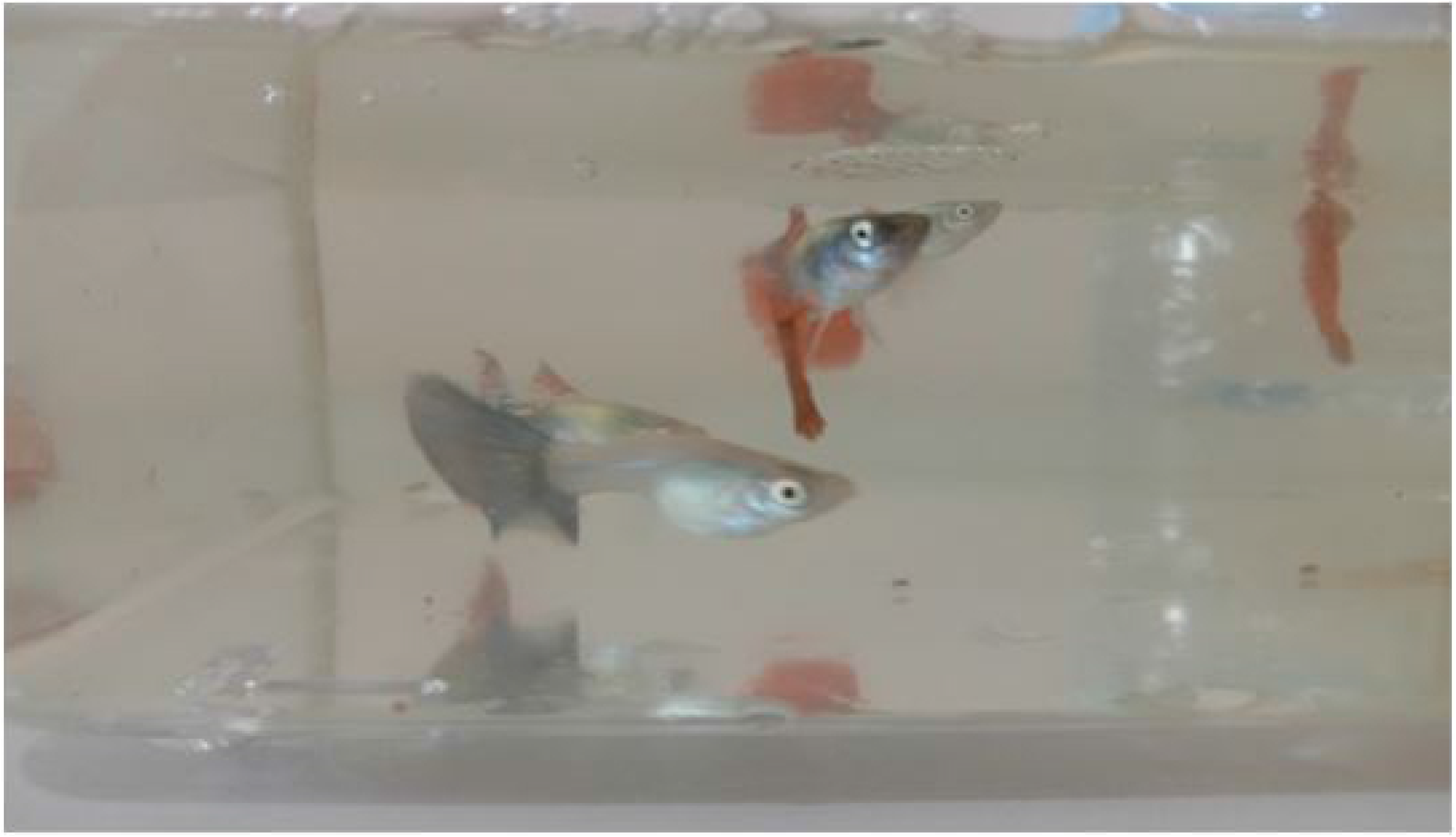
Guppy fish, *Poecilia reticulate* toxicity test with *Clitoria ternatea* flower extracts

**Fig 6.**
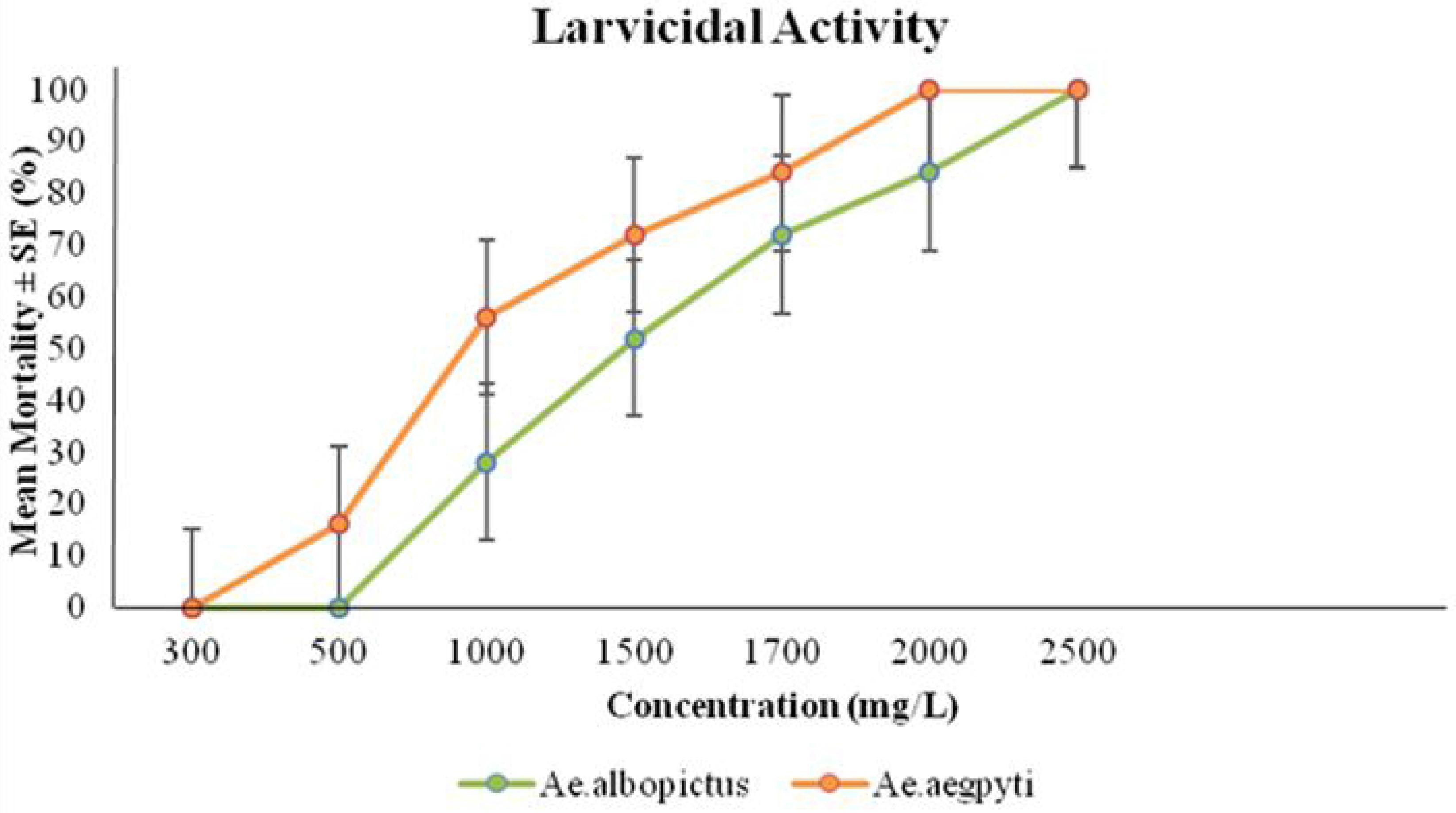
Comparison of larvae mortality rate between *Aedes aegypti* and *Aedes albopictus* for various *Clitoria ternatea* flower extract concentrations

### Morphological view

The morphological view from Fig 7A & Fig 7B are indicating the presence of *C. ternatea* flower extracts in the midgut content by dark greenish colour of extracts.

**Fig 7.**
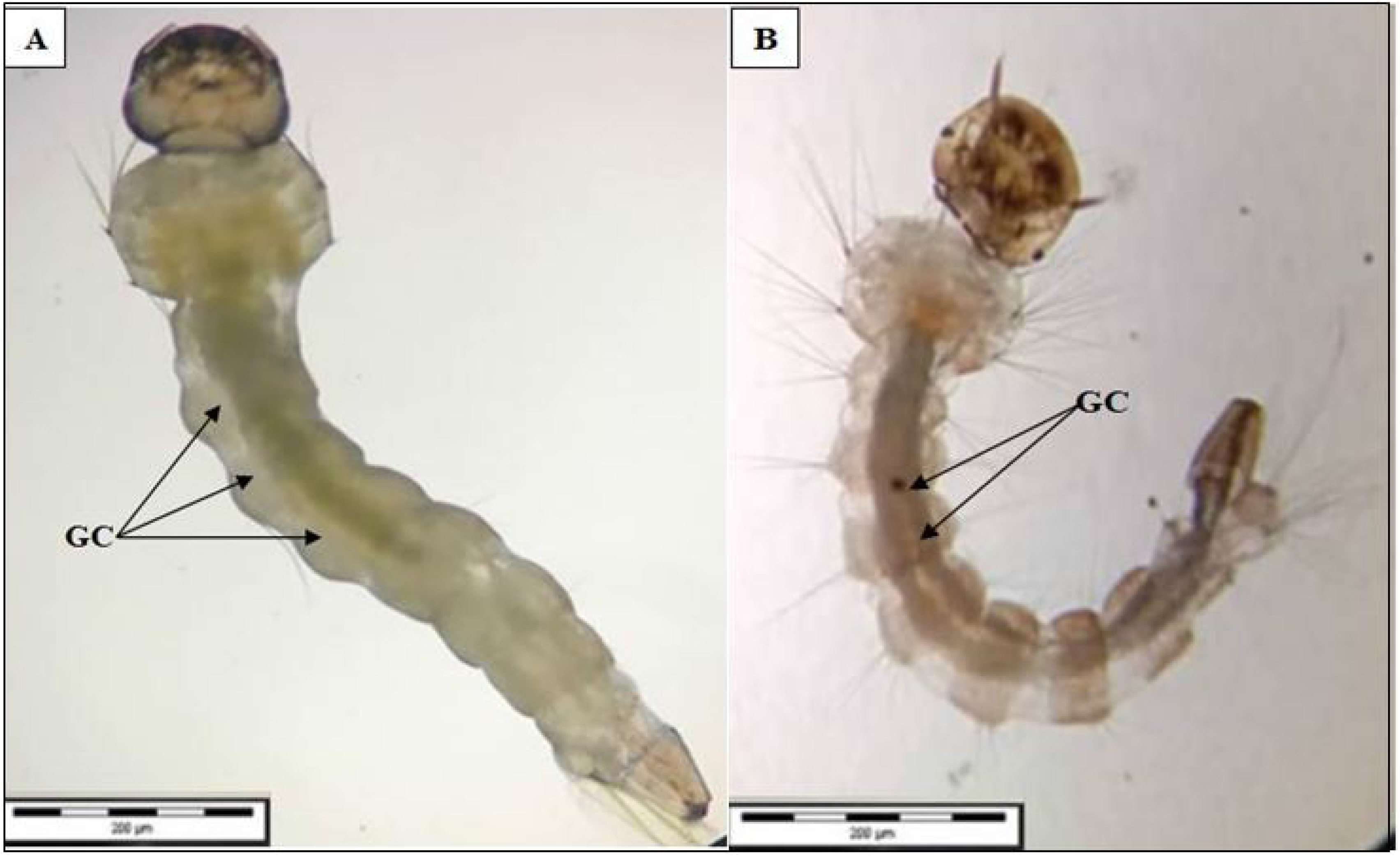
Morphological midgut content induced by *Clitoria ternatea* flower extract (A) Midgut content view in larvae of *Ae. aegypti* (B) Midgut content view in larvae of *Ae. albopictus* Note: Arrows indicating the plant extracts (dark greenish colour), GC: gut content (after 24hours)

## Discussion

Results of this study have shown that phytochemical compounds extracted from *C.ternatea* flower may be innovative and integrated conception of bio insecticidal. Thus this study has set as an alternative control measures against *Aedes* compared to synthetic insecticides. Additionally, *C. ternatea* flower extracts may be more effective than single based active compound due to its active ingredients synergisms which may be effective in managing resistant population of mosquitoes. In this study, we have found total of 16 chemical compounds and 6 chemical compounds have been reported to posse’s direct effects on insecticidal, larvicidal and pupicidal effects. Namely 6 chemicals used for insecticidal properties in previous studies were Glycerin, 2-Hydroxy-gamma-butyrolactone, Neophytadiene, n-Hexadecanoic acid, cis-Vaccenic acid, and Octadecanoic acid with total of 28.7%. All these chemical components has direct effects on the larvicidal activity and there are also many other compounds which may act synergistically in the process of larvicidal activity. Similarly, Ravi et al.[2,4] has also discussed about the synergetic effects of multiple chemical compounds from plant extract against *Aedes* larvae.

Besides that, previous study of Mathew et al.[9] on *Clitoria ternatea* extracts did not show the mechanism of larvicidal activity. However, to elucidate the effects of larvicidal mechanisms, this current study has shown the midgut content ingestion, morphological photogram (Fig 7A and &7B). The ingested plant extract shown in Fig 7A and &7B were similar to previous study done with *Azolla pinnata* plant extracts (green colour) in the larvae midgut content after 24 hours of incubation [4]. Thus, it can be congruently observed that the *C. ternatea* extract ingestion mechanism was used in killing effects of *Aedes* larvae. As the results of *C. ternatea* extract in guts of larvae, the potential commercialization of this extracts would be on liquid based techniques. Previous example of similar possible commercial application like “Dalmatian Powder”, dissolved in liquid for its application[33]. Additionally, the storage condition of the crude extract in this study was at −4°C before its application under room temperature. Similarly Ravi et al.,[4] have prevented the degradation of crude extracts by storing it at −4°C before its application.

The residual activity of *C. ternatea* flower extract showed till 48 hours in water. Moreover, Ullah et al., [34] recorded that 5 different plant species have residual activities against *Culex quinquefasciatus* till 72 hours upon its application. Elsewhere, a study on *Azolla pinnata* extract has suggested that 48 hours of residual activity in water and plant based applications were much more shorter in residual effects compared to synthetic pesticides [4]. Thus, it can be concluded that, plant based insecticides possess lesser residual activity compared to synthetic chemicals. Additionally, the *C. ternatea* plant extracts did not posses any toxic effects on fish. According to Pereira and Olivaira [35], *Poecilia reticulate* has important potential effects to predate *Aedes aegypti* larvae and this will help to eliminate the breeding ground of larvae. Thus, without the *C. ternatea* extracts toxic effects on *Poecilia reticulate*, we could integrate both of the application using *C. ternatea* and fish to control *Aedes* larvae. In addition to that, the toxicity results in this current study were also similar to the previous study based on *Azolla pinnata* plant [2, 4]. Thus, the use of *C. ternatea* extracts is not toxic for the environment and it is safer to be applied compared to chemical application in controlling *Aedes* larvae.

In addition to that, metabolite study of *C. ternatea* extracts showed different types of phenols such as anthocyanin, flavonoid and tannin. Anthocyanin, flavonoid and tannin were present in both methanol and ethanol extraction for *C. ternatea* extracts compared to other extractions (Table 1). Divya et al., [14] reported the similar findings on *Clitoria ternatea* blue and white flowered leaves which recorded highest polyphenol isolation using methanol extraction techniques. Moreover, this was further proven with thin layer chromatography and paper chromatography techniques, whereby all the extracts were found to have three spots upon fixing with iron (iii) chloride, FeCl_3_. Concurrently, Sonam et al., [36] has also reported the same results with medicinal plant using iron (iii) chloride, FeCl_3_ to identify only phenolic compound on chromatography paper. Finally, the antioxidant test for *C. ternatea* flower extracts showed aqueous extract has the highest antioxidant property compared to other extractions. The aqueous extract found to have high antioxidant because water extract can absorb antioxidant compound better and chemicals [37].

Thus, it can be concluded that, *C. ternatea* flower extracts based insecticides may possess lesser residual activity compared to synthetic chemicals, contains multiple chemical compounds which may solve the single based insecticide resistant problems. Finally, although, *C. ternatea* flower extracts may be used as bio-larvicides, future testing would have to be conducted to validate its long term effects on human health and other organisms in environment.

## Conclusion

We can conclude that this current study has shown colossal effectiveness of *Clitoria ternatea* flower extracts against major dengue vectors during the early 4^th^ instar larvae stages. Additionally, current findings showed that crude extracts bioactive molecules to be effective and may be developed as biolarvicides for *Aedes* mosquito vector control programs. Furthermore, this study also provided a baseline understanding for future research work on the field applications of *C. ternatea* flower extracts which could be tested for its long term effects on other non target organisms, including human health.

## Acknowledgments

We would like to thank the Quest International University Internal Funds, Universities staffs and research students who have contributed in this work.

